# A conserved mechanism regulates reversible amyloids *via* pH-sensing regions

**DOI:** 10.1101/2022.03.21.484600

**Authors:** Gea Cereghetti, Vera Maria Kissling, Lisa Maria Koch, Alexandra Arm, Pavel Afanasyev, Miriam Linsenmeier, Cédric Eichmann, Jiangtao Zhou, Yiping Cao, Dorota Maria Pfizenmaier, Sonja Kroschwald, Thomas Wiegand, Riccardo Cadalbert, Daniel Böhringer, Raffaele Mezzenga, Paolo Arosio, Roland Riek, Matthias Peter

**Affiliations:** Institute of Biochemistry, Department of Biology, ETH Zürich, Otto-Stern-Weg 3, 8093 Zürich, Switzerland; Life Science Zürich, PhD Program for Molecular Life Sciences, 8057 Zürich, Switzerland; Life Science Zürich, PhD Program for Biomolecular Structure and Mechanism, 8057 Zürich, Switzerland; Cryo-EM Knowledge Hub (CEMK), ETH Zurich, Otto-Stern-Weg 3, CH-8093 Zürich, Switzerland; Department of Chemistry and Applied Biosciences, Institute for Chemical and Bioengineering, ETH Zurich, Vladimir-Prelog-Weg 1-5/10, 8093, Zurich, Switzerland; Department of Chemistry and Applied Biosciences, Laboratory of Physical Chemistry, ETH Zurich, 8093 Zürich, Switzerland; Department of Health Sciences & Technology, ETH Zürich, Schmelzbergstrasse 9, 8092 Zürich, Switzerland

**Author notes:** To whom correspondence should be addressed: Email addresses. These authors contributed equally to this work (Vera Maria Kissling, Lisa Maria Koch, Alexandra Arm, Thomas Wiegand). Current address: Max-Planck-Institute for Chemical Energy Conversion, Stiftstrasse 34-36, 45470 Mülheim an der Ruhr, Germany, and Institute of Technical and Macromolecular Chemistry, RWTH Aachen University, Worringerweg 1, 52074 Aachen, Germany.

## Abstract

Amyloids were long viewed as irreversible, pathological aggregates, often associated with neurodegenerative diseases^1^. However, recent insights challenge this view, providing evidence that reversible amyloids can form upon stress conditions and fulfil crucial cellular functions^2^. Yet, the molecular mechanisms regulating functional amyloids and the differences to their pathological counterparts remain poorly understood. Here we investigate the conserved principles of amyloid reversibility by studying the essential metabolic enzyme pyruvate kinase (PK) in yeast and human cells. We demonstrate that PK forms stress-dependent reversible amyloids through a pH-sensitive amyloid core. Stress-induced cytosolic acidification promotes aggregate formation *via* protonation of specific glutamate (in yeast) or histidine (in human) residues within the amyloid core. Our work thus unravels a conserved and potentially widespread mechanism underlying amyloid functionality and reversibility, fine-tuned to the respective physiological cellular pH range.

## Main Text

Protein aggregates, and in particular amyloids, are associated with neurodegenerative diseases, such as amyotrophic lateral sclerosis (ALS), Alzheimer’s (AD) and Parkinson’s disease (PD)^1^. However, an expanding number of amyloids have also been found to exert important physiological roles. Indeed, functional amyloids have been described in bacteria, fungi, plants and mammals, fulfilling a wide variety of tasks including long-term memory formation, stress response, metabolism regulation, hormone storage and melanin production^2^. Interestingly, recent structural studies of functional amyloids highlighted their similarity to pathological aggregates, raising the question of what differentiates functional from toxic aggregates^3^. A key difference between the two resides in the presence of cellular mechanisms that govern functional amyloid formation and disassembly, restricting these processes temporally and spatially^4^. However, although functional amyloids exist in all kingdoms of life, only a handful was shown to be reversible in a physiological context, and the exact molecular mechanisms dissolving functional amyloids remain largely unknown.

The yeast pyruvate kinase Cdc19 is an example of an enzyme forming such reversible, functional amyloids. Pyruvate kinase (PK) is essential and regulates a pivotal reaction coupling glucose and energy metabolism. Specifically, it catalyses the last rate-limiting step of glycolysis, converting phosphoenolpyruvate and ADP into pyruvate and ATP. Under favourable growth conditions, Cdc19 is soluble and active as a tetramer in the cytoplasm. Upon stress such as glucose starvation or heat shock, Cdc19 rapidly aggregates into solid cytoplasmic foci (Fig. 1A and^5, 6^). In its aggregated form, Cdc19 is inactive and resistant to stress-induced degradation, while re-solubilization restores its enzymatic activity and energy production^7^. Thus, rapid formation of Cdc19 aggregates shuts down glycolysis and preserves Cdc19 from degradation during stress, while fast aggregate disassembly is crucial to restore energy production and reactivate metabolism. Interestingly, Cdc19 aggregates possess an amyloid structure, both *in vivo* and *in vitro* (Fig. 1A and^5, 7^), and thus resemble irreversible, pathological inclusions found in several neurodegenerative diseases^7, 8^. Yet, Cdc19 amyloids are fully re-solubilized in cells within minutes after stress release.

**Fig. 1.**
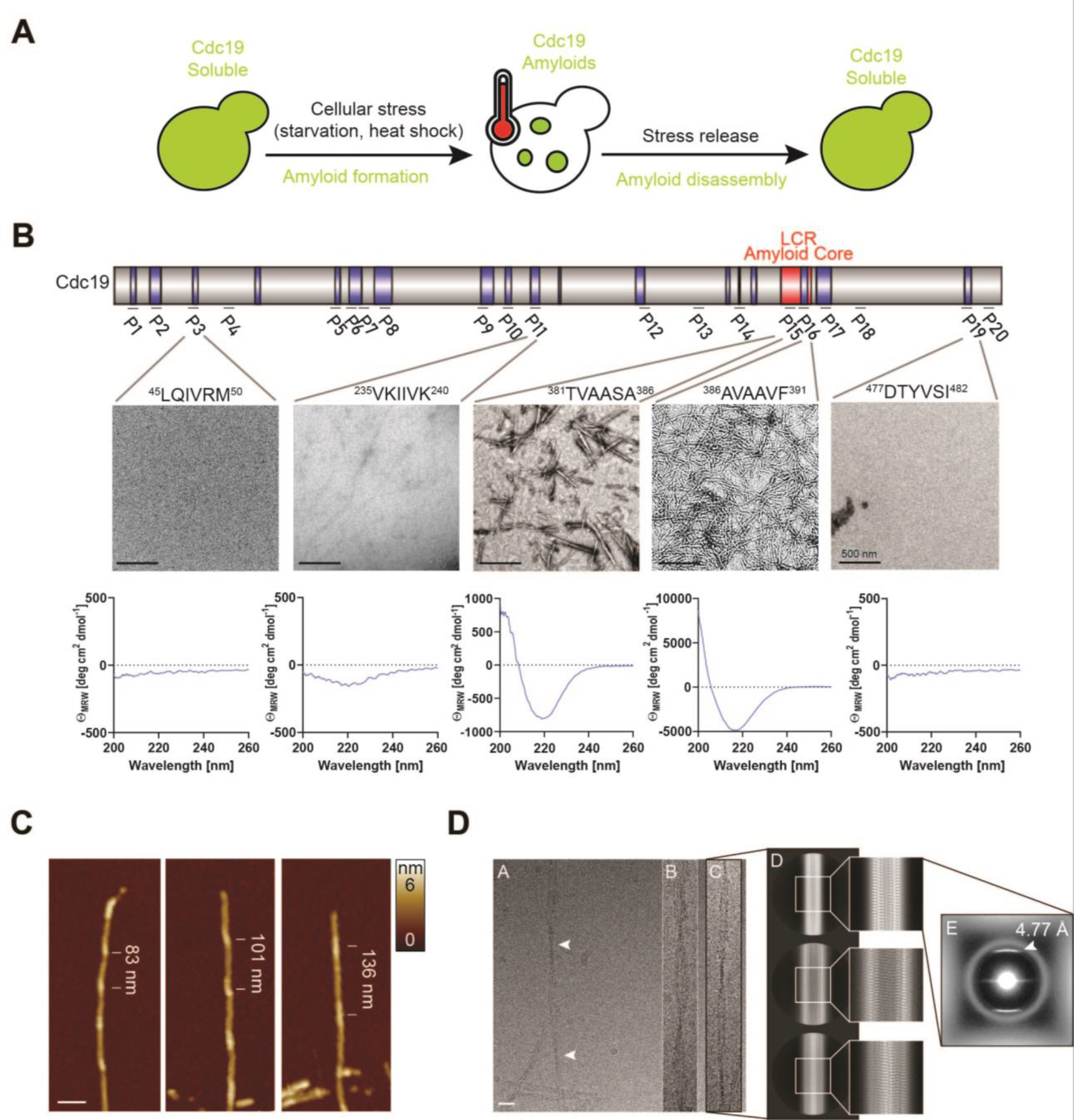
Identification and structural characterization of the amyloid core of Cdc19. (A) Yeast pyruvate kinase (Cdc19) forms reversible, functional amyloids *in vivo* and *in vitro*. Schematic drawing representing Cdc19 localization in yeast cells before, during and after stress. Upon stress Cdc19 forms cytoplasmic aggregates, which have an amyloid structure both *in vivo* and *in vitro*^5, 7^. Aggregation protects Cdc19 from stress-induced degradation and is essential for cell survival to stress. (B) Screening to identify amyloidogenic regions in Cdc19. Schematic representation of the Cdc19 sequence (top). Regions highlighted in blue are predicted by computational tools (ZipperDB^10^ and AmylPred2.0^9^) to be highly amyloidogenic. A predicted aggregation-prone low-complexity region (LCR)^11^ is highlighted in red. 16 hexapeptides (P) corresponding to the regions with highest predicted amyloidogenicity plus 4 negative controls were selected and assessed for their ability to form amyloids by negative staining transmission electron microscopy (TEM) and circular dichroism (CD) spectroscopy. For CD measurements, peptides were fibrilized, collected by centrifugation, washed and measured. Thus, a flat line (as seen for P3 and P19) indicates that no fibrils could be collected. Note that the y-axis is adjusted for each curve for better display. TEM micrographs (image panels) and CD spectra (graph panels) of representative peptides are shown (complete screen results in Extended Data Fig. 1). Please note that the region corresponding to the Cdc19 LCR highlighted in red is by far the most amyloidogenic, and thus defined as the amyloid core. Data are representative of three independent experiments. Scale bar: 500 nm. (C) Structural characterization of the Cdc19 amyloid core by AFM. A synthetic peptide encompassing the amyloid core of Cdc19 (amino acids 376-392) was allowed to form aggregates by incubation for 2 days at 30 °C. Resulting fibrils were analysed by atomic force microscopy (AFM) as described in Materials and Methods, and representative images are shown. The amyloid core of Cdc19 forms fibrils that are polymorphic, similar to published pathological amyloids^46^. Scale bar: 50 nm. (D) Cryo-EM characterization of fibrils formed by the Cdc19 amyloid core. Representative raw micrograph with a clear non-overlapping filament is shown in D^A^. A high level of heterogeneity was identified in this dataset: fibrils of various thickness and cross-over distances (D^A^: 1450 Å – indicated by white arrows, D^B^: 1100 Å; D^C^: 1400 Å), indicating the presence of polymorphism (D^A-C^). All images are raw and to scale (scale bar: 200 Å). (D^D^) 2D-classification of manually selected particles, corresponding to thin fibrils (like in D^A^ and D^B^) revealed class-averages with a rise distance of 4.77 characteristic for separation etween a β-strands in amyloids. (D^E^) power spectrum of one of the 2D class-averages (average of the amplitudes of the particles, contributing to this class).

Here, we exploit pyruvate kinase in yeast and human cells to dissect the molecular mechanisms and structural characteristics underlying amyloid reversibility. Understanding the principles governing amyloid reversibility may not only reveal fundamental insights into this physiological process, but could also have important implications for developing novel strategies to treat amyloid-related diseases.

### Identification and structural characterization of yeast pyruvate kinase (Cdc19) amyloid core

In order to understand the mechanisms regulating reversible amyloid formation of Cdc19, we first sought to identify sequence elements in Cdc19 responsible for its reversible aggregation. Using computational tools (AmylPred2.0^9^ and ZipperDB^10^) predicting the propensity of a given sequence to form cross-β structures, we identified several putative amyloidogenic regions (Fig. 1B, upper panel, in blue). A series of 20 hexapeptides distributed along the Cdc19 sequence were then synthesized, 16 of which are in the predicted highly amyloidogenic regions and 4 for control in low amyloidogenic regions. These peptides were then screened for their ability to form amyloids using the amyloid-binding dyes Thioflavin T (ThT) and Congo Red (CR) (Extended Data Fig. 1A and B). ThT- and/or CR-positive peptides were subsequently visualized by negative staining transmission electron microscopy (TEM) (Fig. 1B, middle panels, and Extended Data Fig. 1C) and structurally analysed by circular dichroism (CD) spectroscopy (Fig. 1B, lower panels). These analyses identified a prominent region of interest around peptides 15 and 16, which efficiently formed amyloid fibrils (Fig. 1B). Peptides 11 and 19 yielded amyloid fibrils, however to a much lower degree, as observed by TEM or dye staining, respectively (Fig. 1B, Extended Data Fig. 1A and B). Additional computational analysis using prediction tools to identify aggregation-prone low-complexity regions (LCRs) (e.g. SEG algorithm^11^) identified a single LCR within Cdc19 (amino acids 376-392, Fig. 1B, upper panel, in red)^5^. This sequence has an α-helical structure in tetrameric Cdc19 and contains the experimentally identified amyloidogenic peptides 15 and 16, and is thus likely the main driver of Cdc19 aggregation. Therefore, we termed it “amyloid core” (Fig. 1B, upper panel, in red).

To investigate the defining principles distinguishing Cdc19 amyloids from their pathological counterparts, we further characterized the structural and biophysical features of the Cdc19 amyloid core. Amyloids are characterized by a highly ordered β-sheet-rich structure, where individual β-strands align perpendicularly to the fi ril axis with a spacing of 4.7 etween adjacent β-strands^12, 13^. Amyloid fibrils are often formed by two or more twisting protofilaments^14, 15^, which create rather regular crossovers that can be readily observed in TEM or atomic force microscopy (AFM)^16^. Indeed, characterizing fibrils formed by the Cdc19 amyloid core (sequence 376-392) by AFM revealed different unbranched, left-hand twisted fibrillar structures with a diameter of 23 ± 4 Å and a periodicity (i.e. crossover distance) of 1010 ± 180 Å (Fig. 1C and Extended Data Table 1), resembling classical amyloid fibrils. These findings were complemented by cryoEM analysis of the Cdc19 amyloid core (Fig. 1D), which showed a characteristic staggered β-sheet repeat structure of the fibrils in 2D analysis with a rise distance of 4.77 (Fig. 1D, white arrow). oreover, the cryo-micrographs corroborated the co-existence of different fibril structures with varying crossover distances (up to approximately 2000 Å), protofilament numbers and fibril thicknesses (Fig. 1D). Interestingly, such a high degree of structural polymorphism^17^ is usually associated with pathological amyloids like α-synuclein or amyloid-β^3, 13^. However, despite these remarkable structural similarities (Extended Data Table 1), Cdc19 amyloids in cells can efficiently form and disassemble under physiological conditions. Thus, amyloid reversibility is likely governed by defined cellular activities and/or specific amino acid sequences and their biochemical properties.

### Cdc19 amyloid reversibility depends on pH-sensing glutamic acids in its amyloid core

Irreversible, pathological amyloids are often characterized by the presence of large hydrophobic interfaces in their core^13, 18^, while aggregates that can be disassembled and re- solubilized in a physiological context contain more hydrophilic residues, as found in LCRs and prion-like domains^19, 20^. Indeed, reversible aggregates of hnRNPA2-LCR or FUS-LCR are almost completely devoid of non-polar residues like alanine, valine, isoleucine and leucine, but are rich in asparagine and glutamine as well as phosphorylatable residues (Extended Data Table 2)^20^. Mutational analysis and mutations found in ALS families^21^ confirmed that the asparagine residues in the core of hnRNPA2 are essential for reversibility. Similarly, the core of the functional amyloid Orb2 is hydrophilic, owing to 7 histidines and 20 glutamines out of 31 residues^22^. Interestingly, the amyloid core of Cdc19 is mainly formed by hydrophobic amino acids (Fig. 2A), but also contains multiple serine and threonine residues, whose phosphorylation has been shown to regulate Cdc19 aggregation *in vivo*^5^. In addition, the Cdc19 amyloid core contains charged residues (Fig. 2A), which confer a different set of chemical properties, including the ability to sense and react to pH changes. Since reduction in intracellular pH is a conserved signal regulating many cellular processes in response to nutrient starvation and other stress conditions^23–25^, we investigated if the formation and disassembly of Cdc19 amyloids could be influenced by pH.

**Fig. 2.**
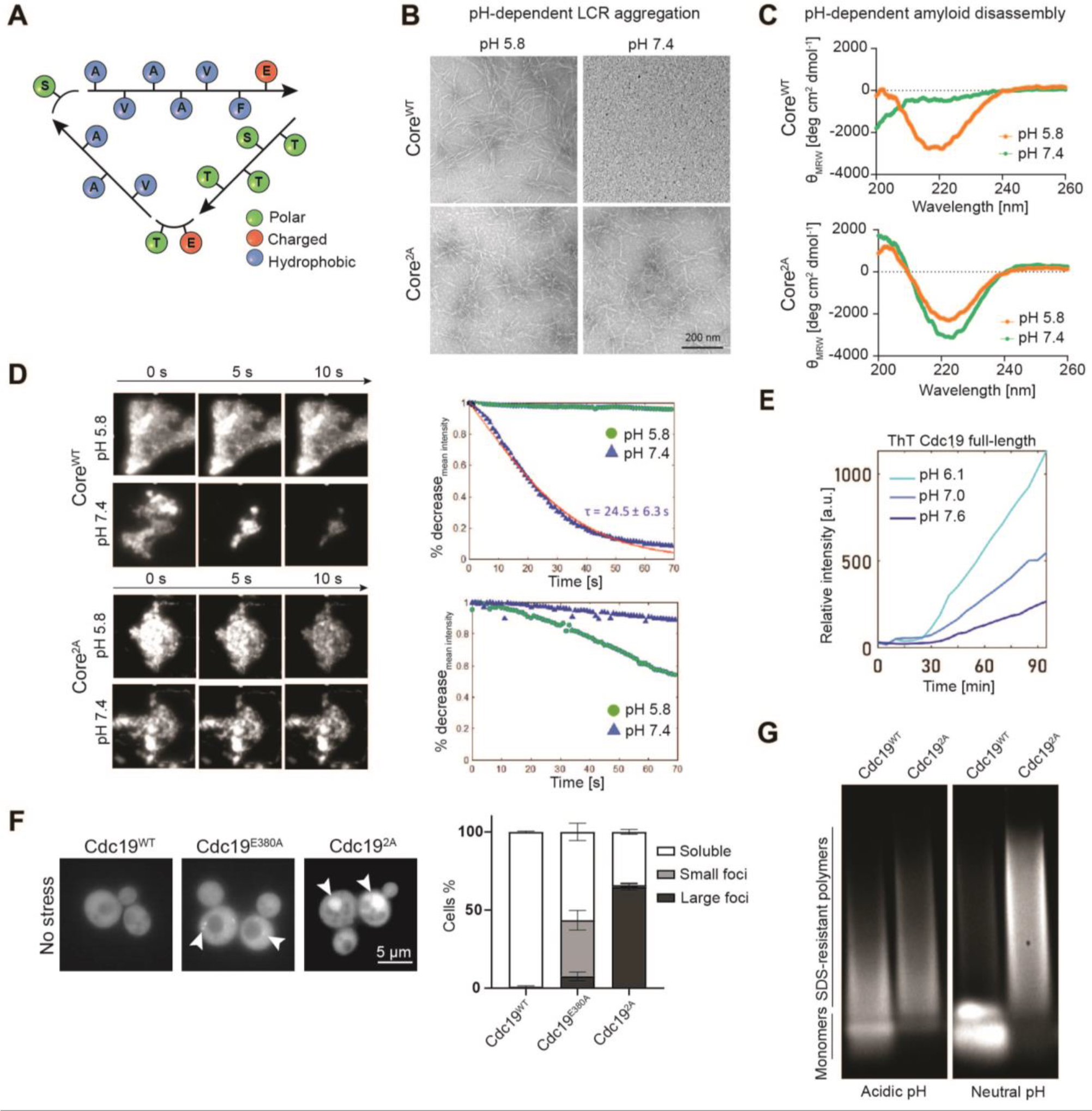
Formation and disassembly of Cdc19 amyloids is regulated by pH, *via* protonation of two specific glutamic acid residues (E380 and E392) in the amyloid core. (A) Cdc19 amyloid core sequence. The sequence of the Cdc19 amyloid core is represented, and the biophysical characteristics of the different amino acids are indicated by different colours. (B) Cdc19 amyloid core forms amyloid fibrils only at physiologically low pH. Wild-type (Core^WT^) or mutant (Core^2A^, mutations: E380A, E392A) Cdc19 amyloid cores were incubated at pH 5.8 or pH 7.4 for two days and imaged by negative staining TEM. Note that Core^WT^ forms fibrillar aggregates at physiologically low pH corresponding to the intracellular pH of stressed cells, while it remains soluble at neutral pH corresponding to the intracellular pH of growing cells. Core^2A^ peptides are pH-insensitive and form fibrils under both conditions. n = 3. Scale bar: 200 nm. (C) Pre-formed amyloids are rapidly dissolved at neutral pH corresponding to the intracellular pH of growing cells. Core^WT^ and Core^2A^ peptides were allowed to form fibrils overnight at pH 5.8, and their secondary structure was determined by circular dichroism (CD) spectroscopy (orange). CD spectra were re-measured after increasing the pH to 7.4 (green). Note that upon pH increase Core^WT^ transitions from a β-sheet-rich, amyloid structure to a random coil structure, while Core^2A^ maintains its β-sheet-rich signature. n = 3. (D) pH-mediated amyloid disassembly occurs within seconds. Wild-type (Core^WT^, upper panels) or the pH-insensitive mutant (Core^2A^, lower panels) amyloid cores of Cdc19 were allowed to aggregate overnight at pH 5.8. Fibrils were then stained with Thioflavin T (ThT), trapped in a microfluidic device, and flushed with buffers at pH 5.8 or pH 7.4 as indicated. Aggregates were imaged over time by fluorescence microscopy and representative images of three independent experiments are shown (image panels). Aggregate disassembly was quantified as percentage (%) decrease in mean fluorescence intensity over time (graphs). Characteristic time of aggregate disassembly is reported as τ. Note that Core^WT^ amyloids disassembled within a few seconds at pH 7.4, while Core^2A^ amyloids remained stable under these conditions. (E) pH affects aggregation of full-length Cdc19. Purified full-length Cdc19^WT^ was allowed to aggregate at different pH conditions, and aggregation kinetics were monitored by ThT fluorescence. Lower pH strongly accelerated Cdc19 aggregation (n = 3). (F) Aberrant pH-sensing leads to constitutive aggregates *in vivo*. Cells expressing GFP- tagged Cdc19^WT^, Cdc19^E380A^ or Cdc19^2A^ were grown in SD-full media and imaged by fluorescence microscopy. Arrows mark aggregates. Note that introducing one (E380A) or both (E380A, E392A) pH-insensitive mutations leads to constitutive aggregates independent of intracellular pH. The intensity of the image showing Cdc19^2A^ was adjusted for etter visualization. Scale ar: 5 μm. The percentage of cells with solu le Cdc19 or bearing small or large aggregates was quantified by manual counting. Graph represents mean ± SEM (n = 3, at least 50 cells were counted for each condition). (G) Maintaining neutral pH during starvation prevents Cdc19 amyloid formation *in vivo*. Cells expressing GFP-tagged wild-type (Cdc19^WT^) or the pH-insensitive Cdc19 mutant (Cdc19^2A^) were grown to stationary phase in SD-full media. The media was either kept at its normal pH of around 5, or adjusted to pH 7.5, and cells were starved for additional 8 days. The presence of starvation-induced amyloids was analysed by SDD-AGE using an α-GFP antibody. Note that at low pH Cdc19^WT^ develops SDS-resistant structures indicative of amyloids, while this is prevented at neutral pH conditions. In contrast, Cdc19^2A^ assembles SDS-resistant amyloids independent of pH (n = 3).

Strikingly, we observed that the Cdc19 amyloid core (Core^WT^) forms fibrillar aggregates at physiologically low pH (pH 6, corresponding to the intracellular pH of starved or heat shocked yeast cells^26^), while it remains soluble at neutral pH (corresponding to the pH of growing cells^26^) (Fig. 2B, upper panels). Mutational analysis confirmed that these pH changes are sensed by the two protonatable glutamic acid residues within the amyloid core. Indeed, substituting both glutamic acids with non-charged alanine residues (E380A, E392A; Core^2A^) mimicking the neutral charge of protonated glutamic acids, or un-charged polar residues (E380Q, E392Q; Core^2Q^) led to pH-insensitive, constitutively aggregating amyloid cores (Fig. 2B, lower panels, Extended Data Fig. 2A). The uncharged alanine or glutamine residues mimic the neutral charge of protonated glutamic acids present at low pH, thus resulting in constitutive aggregates even at neutral pH. Importantly, neutral pH could readily disassemble pre-formed Core^WT^ fibrils, as shown by CD spectra (Fig. 2C). When Core^WT^ and Core^2A^ peptides were incubated at low pH, oth peptides adopted a typical β-sheet-rich amyloid signature. In contrast to Core^2A^ fibrils, increasing the pH from 5.8 to 7.4 led to the rapid disappearance of Core^WT^ fibrils, which adopted a random coil structure. Solid-state NMR measurements exploring the pH-sensitivity of the ^13^C chemical-shift value of the carboxyl carbon^27^ confirmed the glutamic acid residues in the Core^WT^ fibril structure are protonated and thus non-charged, while they become partially deprotonated and thus negatively charged when the pH is increased above pH 6 (Extended Data Fig. 2B). To quantify the kinetics of the Cdc19 amyloid core disassembly, we imaged Core^WT^ and Core^2A^ aggregate dissolution in real-time in a microfluidic chamber upon switch of the medium pH from 5.8 to 7.4 (Fig. 2D). Even very large Core^WT^ amyloid aggregates tens of μm in diameter dissolved with a half-time of less than 25 seconds, in strong contrast to Core^2A^ or pathological amyloids, which in physiological contexts are usually stable and irreversible^4^. We conclude that physiological pH changes regulate rapid Cdc19 amyloid fibril formation and disassembly *via* reversible protonation of E380 and E392 controlling electrostatic repulsion.

We next tested whether pH-regulation of the amyloid core also affects reversible aggregation of full-length Cdc19 *in vitro* and *in vivo*. Lower pH facilitated aggregation of purified Cdc19 as measured by ThT fluorescence, while higher pH slowed aggregation (Fig. 2E). As expected, the pH-insensitive full-length Cdc19 mutants carrying one (*cdc19-E380A*) or both pH-insensitive mutations (*cdc19-E380A, E392A*, henceforth called 2A) were extremely aggregation-prone and rapidly formed large oligomers and fibrils even at neutral pH (Extended Data Fig. 2C and D, arrows). Interestingly, yeast cells expressing the pH-insensitive Cdc19^2A^ mutant fused to GFP from the endogenous locus were strongly impaired for growth and accumulated large foci even in no stress conditions (Fig. 2F and Extended Data Fig. 2E), while cells bearing Cdc19^E380A^-GFP presented an intermediate phenotype. Both Cdc19^E380A^ and Cdc19^2A^ had reduced protein levels compared to Cdc19^WT^ controls (Extended Data Fig. 2F), suggesting that cells may try to degrade irreversible aggregates.

In the cellular context, physiological cytosolic acidification is a common response to several stresses, such as nutrient starvation or heat shock^25, 26, 28, 29^. Yeast cells usually grow in media with a pH of around 4-5, but they maintain a neutral cytosolic pH under favourable growth conditions. However, stresses such as glucose starvation or heat shock cause a rapid cytosolic acidification, leading to an intracellular pH of around 6^26, 28^. We observed that when yeast cells are exposed to starvation conditions, endogenously expressed wild-type and Cdc19^2A^ formed SDS-resistant amyloids *in vivo*. Strikingly, however, preventing starvation-induced intracellular acidification by switching cells into neutral pH 7.5-adjusted media^25^ was sufficient to abolish the formation of SDS-resistant amyloids for wild-type but not Cdc19^2A^ (Fig. 2G). Thus, stress-induced cytosolic acidification is essential for Cdc19 amyloid formation *in vivo*.

Taken together, these data suggest that the formation and dissolution of Cdc19 amyloids *in vivo* is regulated by cytosolic pH, *via* protonation of two specific glutamic acid residues in the amyloid core. Low pH and consequent glutamic acid protonation result in a non-charged amyloid core, which drives folding into polymorphic amyloid fibrils. Upon return to neutral pH, deprotonation of these glutamic acids and the resulting electrostatic repulsion likely destabilize the core and trigger amyloid dissolution.

### Human pyruvate kinase (PKM2) forms reversible amyloids in a pH-dependent manner

Stress-induced acidification is conserved in eukaryotes, from yeast to plants, insects and mammals^25^. Given the high functional and structural conservation of pyruvate kinase (PK), we next investigated whether pH-dependent regulation of reversible PK amyloids is preserved in human cells. In humans, PK is encoded by two genes that produce a total of four isoforms: PKL and PKR, which are expressed only in few cell types, and PKM1 and PKM2, which are ubiquitously expressed in different types of cells and tissues^30^. To examine whether human PKs form pH-regulated reversible amyloids, we subjected RPE-1 cells to starvation, a physiological stress that leads to rapid cytosolic acidification^24, 26^. Interestingly, while PKM1 and PKM2 were soluble and uniformly distributed in the cytoplasm in the absence of stress, PKM2 but not PKM1 formed cytosolic foci upon nutrient starvation in which glucose (Glc) and growth factors (fetal calf serum, FCS) were removed (Fig. 3A). Similarly, artificially reducing cytosolic pH by treating cells with dimethyl amiloride (DMA) or siRNA-mediated depletion of the sodium- hydrogen exchanger 1 (NHE1)^24^, also triggered PKM2 aggregation (Extended Data Fig. 3A and B). NHE1 is the most abundant sodium-hydrogen exchanger at the plasma membrane^24^ and its knockdown or pharmacological inhibition by DMA was previously shown to strongly reduce cytosolic pH^24^. Stress release by FCS and Glc re-addition to starved cells re-established neutral pH, and rapidly re-solubilized PKM2 aggregates, even in the presence of cycloheximide (CHX), which prevents *de novo* protein synthesis (Fig. 3A, Recovery and Recovery + CHX). In contrast, PKM2 foci persisted when cells were released from starvation into media with FCS/Glc in the presence of DMA, which was shown to maintain low cytosolic pH^24^, demonstrating that high cytosolic pH levels are required for PKM2 aggregate disassembly (Fig. 3A, Recovery + DMA). These data suggest that PKM2 but not PKM1 forms reversible aggregates *in vivo* upon starvation *via* a pH-dependent mechanism.

**Fig. 3.**
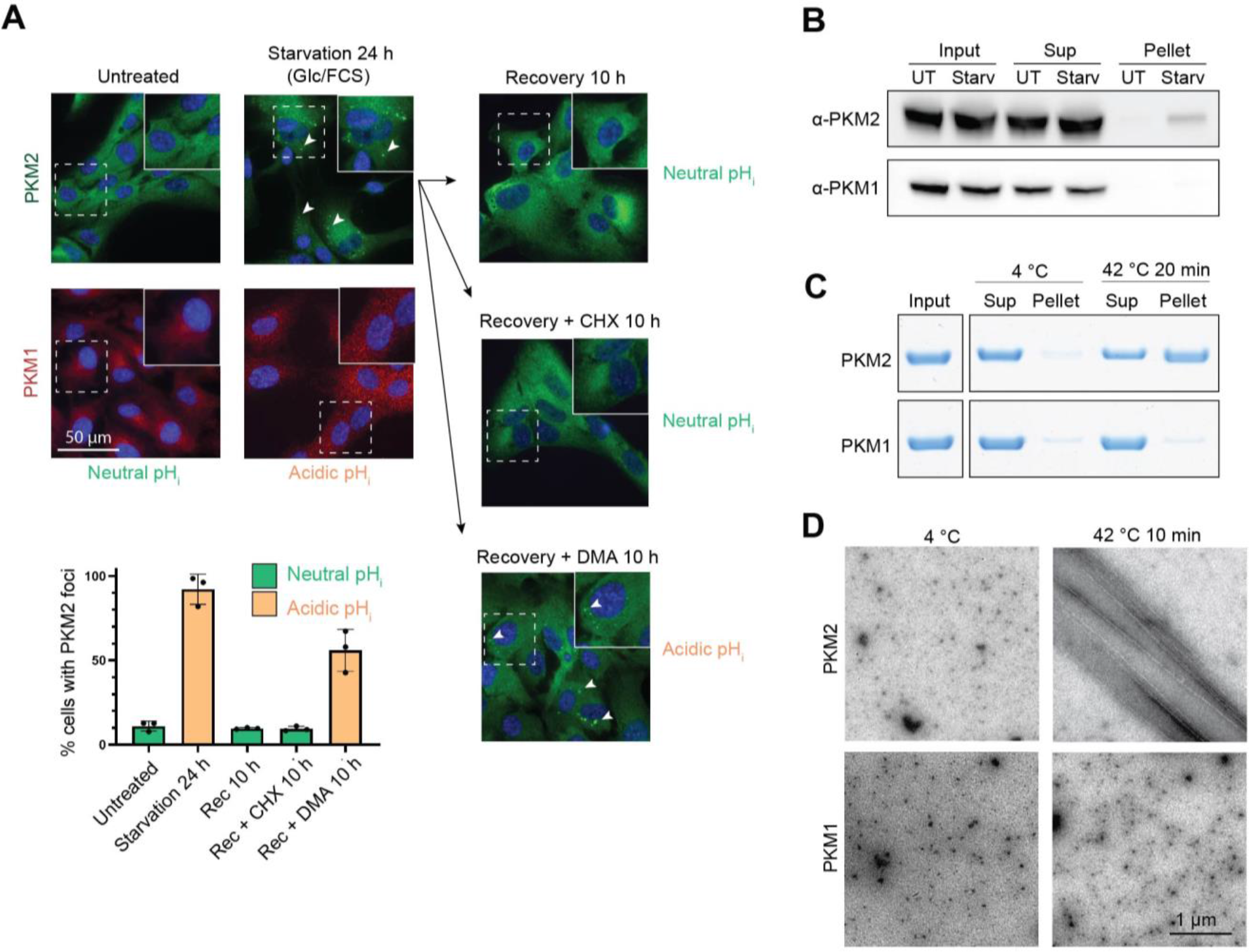
Human pyruvate kinase (PKM2) forms pH-dependent reversible amyloids. (A) PKM2 but not PKM1 forms reversible aggregates in RPE-1 cells upon starvation-induced intracellular acidification. PKM2 and PKM1 localization was analysed in RPE-1 cells by immunofluorescence before, during, and after starvation by removing glucose (Glc) and fetal calf serum (FCS) at the indicated time points. Where indicated, cycloheximide (CHX) or the pH-lowering drug dimethyl amiloride (DMA) were added during recovery to prevent *de novo* protein synthesis or maintain low cytosolic pH, respectively. Scalear: 50 μm. Representative areas (dashed squares) were enlarged x1.5 for better visualization of foci (inserts). The percentage (%) of cells with cytoplasmic PKM2 foci was quantified under the different conditions and indicated as mean ± SEM (n = 3, at least 50 cells were counted for each condition). (B) *In vivo*-formed PKM2 aggregates are insoluble. Extracts of untreated (UT) or Glc/FCS starved (Starv) RPE-1 cells were centrifuged to separate soluble (Sup) and insoluble (Pellet) fractions. Input and a fraction of the soluble (Sup) and insoluble samples were analysed by Western blot with the indicated antibodies (n = 3). (C) Purified full-length PKM2 forms pelletable aggregates upon stress *in vitro*, while PKM1 does not. Purified full-length PKM1 and PKM2 were kept at 4 °C or subjected to heat stress (42 °C, 10 min), leading to mild acidification (around pH 6) of the Tris-based buffer in which the protein is dissolved. Resulting aggregates (Pellet) were separated from soluble protein (Sup) by centrifugation, and a fraction of the supernatant and pellet were analysed by SDS-PAGE and Coomassie blue staining (n = 3). (D) Purified full-length PKM2 forms amyloids upon stress *in vitro.* Purified full-length PKM2 and PKM1 were visualized by negative staining TEM before or after heat stress (42 °C, 10 min). Note that upon heat stress-induced mild acidification, PKM2 forms amyloid-like filaments *in vitro* similar to pathological aggregates, while PKM1 remains solu le. Scale ar: 1 μm.

Starvation-induced PKM2 foci could be pelleted from RPE-1 cell lysates by centrifugation (Fig. 3B), indicating a rather stable structure. To demonstrate that the different aggregation behaviour is intrinsic to the specific PKM isoform, we recombinantly expressed and purified PKM1 and PKM2. Both purified full-length proteins were soluble at 4 °C. However, upon reduction of pH from 7.4 to ∼ 6 triggered by a mild heat shock^31^ (Fig. 3C) or buffer exchange (Extended Data Fig. 3C), PKM2 but not PKM1 formed large, stable assemblies that could be pelleted by centrifugation. TEM analysis revealed PKM2 fibrils with a characteristic amyloid morphology, while PKM1 remained soluble under these conditions (Fig. 3D). Indeed, PKM2 was previously shown to precipitate with beta-isox^32^, a compound known to bind amyloidogenic proteins^6, 33^, and was stained *in vitro* by ThT^34^, further confirming the amyloid structure of these aggregates. While PKM1 and PKM2 were enzymatically active in their soluble state, PKM2 was rapidly inactivated upon fibril formation (Extended Data Fig. 3D). We conclude that, similar to the mechanism described for Cdc19 in yeast, stress-induced changes in cytosolic pH trigger the formation of reversible PKM2 amyloid-like aggregates in human cells. We speculate that this physiological mechanism inactivates and protects PKM2 from stress-induced degradation, owing to the high protease resistance of the amyloid fold^35^.

### Reversibility of PKM2 aggregates depends on a pH-sensing histidine in its amyloid core

To dissect the molecular mechanisms underlying the differential behaviour of PKM2 and its non-aggregating isoform, PKM1, we compared their primary sequence. The two isoforms are produced by alternative splicing of the PKM gene and their sequence is nearly identical (96% sequence identity). The exception is a short region (Fig. 4A, between dashed lines) encoded by exon 9 in PKM1 and thus absent in PKM2, and encoded by exon 10 in PKM2 though excluded in PKM1. Computational analysis of the PKM2 protein sequence with AmylPred2.0^9^ and SEG^11^ predicts an amyloid-prone LCR located in the first half of the PKM2-specific exon 10 (Fig. 4A, highlighted in red). In contrast, exon 9 unique to PKM1 displays neither amyloidogenic propensity nor an LCR. Interestingly, while the putative PKM2 amyloid core is located in the same protein region as the amyloid core in Cdc19, the yeast and human sequences are vastly different (Fig. 4A compared to Fig. 2A). To test whether this putative amyloid core in PKM2 was indeed sufficient to form reversible aggregates, we fused GFP to the predicted PKM2 amyloid core (residues 372-402) or to the corresponding region of PKM1, and expressed these fusion proteins in yeast cells (Fig. 4B). While under exponential growth conditions GFP was soluble and dispersed in the cytoplasm, upon stress the PKM2 amyloid core triggered the formation of reversible GFP aggregates which were readily re-solubilized upon stress release. In contrast, the PKM1 region did not trigger GFP aggregation under these conditions. These results demonstrate that the ability to reversibly aggregate is intrinsic to the PKM2 amyloid core, and does not require human-specific factors.

**Fig. 4.**
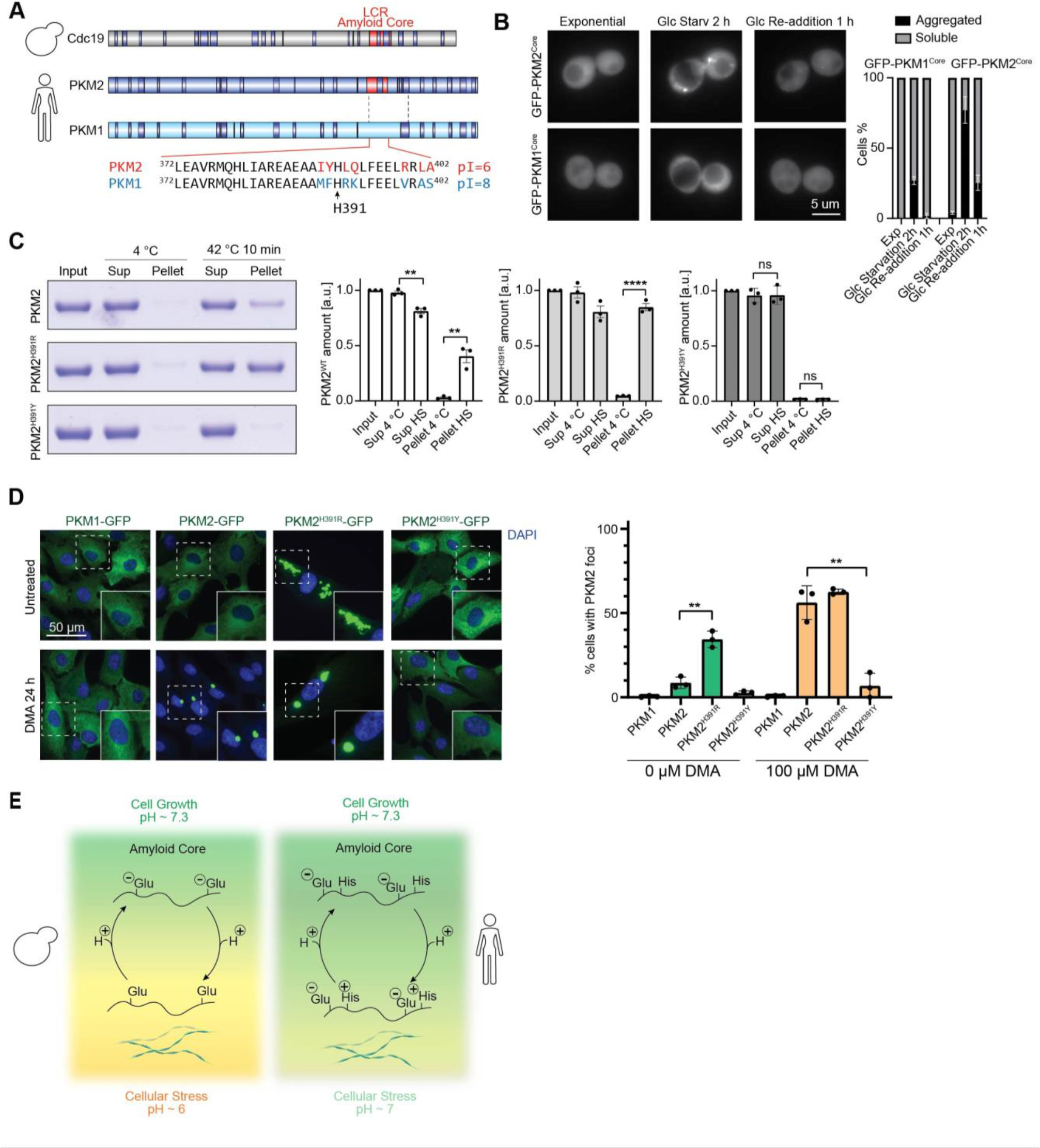
pH-sensing amyloid cores are evolutionarily conserved “amyloid on/off switches”. (A) Schematic representation of yeast Cdc19 and the human pyruvate kinase homologues PKM1 and PKM2. PKM1 and PKM2 are produced from a single gene by alternative splicing and most of their sequence is identical, except for a short region located between the two dashed lines. Regions with amyloidogenic properties predicted by AmylPred2.0 are highlighted in dark blue. The region with highest amyloidogenicity in Cdc19 and PKM2 overlaps with a predicted low-complexity region (LCR) highlighted in red, which is absent in PKM1. The amino acid sequence of this region is listed for PKM2 and PKM1, and different amino acids are highlighted in red and blue, respectively. The theoretical isoelectric point (pI) of these sequences is indicated. (B) The LCR of PKM2 is sufficient to induce reversible aggregation of an otherwise soluble protein *in vivo*. The putative amyloid core of PKM2 or its corresponding region in PKM1 were fused to GFP and expressed in yeast. Cells were then imaged before, during and after starvation at the indicated time points. Percentage (%) of cells with GFP-aggregates is indicated in the graph as mean ± SEM (n = 3, at least 50 cells per time-point per condition were quantified). Scale ar: 5 μm. (C) Mutations of H391 result in hyper-aggregating or non-aggregating full-length PKM2. H391 in full-length PKM2 was mutated to arginine to mimic a positively charged histidine or to tyrosine to mimic a mutation found in Bloom syndrome patients. Wild- type or mutant PKM2 proteins were purified and either kept at 4 °C or subjected to a pH-lowering heat stress (42 °C, 10 min). The resulting insoluble aggregates were separated from soluble protein by centrifugation, and a fraction of the supernatant (Sup, containing soluble protein) and the pellet (Pellet, containing aggregates) were analysed by SDS-PAGE and Coomassie blue staining. Band intensity was quantified using ImageJ and shown as mean ± S.E.M in the bar graphs (n = 3, two-tailed Student’s t-test, *\*\*P_WT,Pellet_* = 0.0031, *\*\*P_WT,Sup_* = 0.004, *\*\*\*\*P_H391R,Pellet_* < 0.0001). Note that the H391R mutation results in a hyper-aggregating PKM2 mutant, while the H391Y mutation abrogates PKM2 aggregation. (D) H391 senses cytosolic pH and regulates PKM2 amyloid formation *in vivo*. GFP- tagged PKM2, PKM2^H391R^ and PKM2^H391Y^ were overexpressed in RPE-1 cells and imaged by fluorescence microscopy in untreated cells or cells treated with the pH- lowering drug D A (100 μ) for 24 h. Note that PK 2 forms pH-dependent aggregates, while the low pH-mimicking PKM2^H391R^ mutant constitutively aggregates independently of cytosolic pH. In contrast, PKM2H^H391Y^ does not aggregate upon stress even if overexpressed. Data are representative of three independent experiments. Scale ar: 50 μm. Representative areas (dashed squares) were enlarged x1.5 for better visualization of foci (inserts). The percentage (%) of cells with cytoplasmic PKM2 foci was quantified under different conditions and is indicated in the graph as mean ± SEM (n = 3, at least 50 cells were analysed for each condition, two-tailed Student’s t-test, *\*\*P_WT-H391R_* = 0.0017, *\*\*P_WT-H391Y_* = 0.0023). (E) The basic principle of pyruvate kinase amyloid reversibility is conserved from yeast to humans. Reversible amyloids of pyruvate kinase are regulated by pH-sensing amyloid cores, which use protonatable residues (glutamic acids in yeast, and histidine in human) to sense stress-induced changes in intracellular pH. Their protonation results in a net charge of 0 of the amyloid cores, which then folds into β-sheet-rich structures and triggers the formation of amyloids. Deprotonation upon return to a neutral pH causes electrostatic repulsion, allowing amyloid re-solubilization. The pH- sensing residues have been adapted through evolution to respond to the stress-induced cytoplasmic pH changes that are characteristic for different organisms.

Interestingly, the PKM2 amyloid core sequence differs from PKM1 by only 6 residues (Fig. 4A), four of them located around Histidine-391 (H391). Due to their pK_a_ of 6, histidines have previously been suggested to function as pH-sensors in mammalian cells^36^, and we thus hypothesized that H391 protonation regulates reversible PKM2 aggregation. Specifically, stress-induced acidification of the cytoplasm may lead to protonation of H391, which analogous to Cdc19 would result in a net charge of 0 in the amyloid core, allowing the formation of reversible PKM2 aggregates. By contrast, the different chemical environment surrounding this histidine in PKM1, particularly the presence of two adjacent positively charged residues (Arg-392 and Lys- 393), may prevent PKM1 aggregation. To test the role of H391 for pH-dependent PKM2 aggregation, we compared wild-type with a mutant peptide of the PKM2 amyloid core centre (amino acids 382-402), where H391 is substituted by a positively charged arginine residue, to mimic the expected protonation state in low pH conditions. As expected, while Core^WT^ amyloids rapidly disassembled at high pH, the Core^H391R^ mutant formed insoluble amyloid aggregates resistant to pH changes, implying that H391 is indeed responsible for pH-dependent reversible aggregation of the PKM2 amyloid core (Extended Data Fig. 4A and 4B). To determine the relevance of H391 in controlling aggregation of full-length PKM2 *in vitro*, we purified wild-type and PKM2 mutant proteins and analysed their aggregation by pelleting assays after mild heat-shock conditions. Indeed, PKM2^H391R^ proved to be more aggregation-prone than PKM2^WT^ (Fig. 4C). Interestingly, a substitution of H391 to tyrosine (H391Y) has previously been reported in patients affected by Bloom syndrome, a genetic disease characterized by genomic instability and predisposition to cancer development^37^. Strikingly, purified PKM2^H391Y^ was unable to form aggregates under these conditions (Fig. 4C), supporting the notion that the positive charge of protonated H391 is required to promote amyloid formation of full-length PKM2.

Finally, we investigated whether pH-dependent PKM2 aggregation *in vivo* also depends on H391 in the amyloid core. GFP-tagged PKM2, PKM2^H391R^ and PKM2^H391Y^, and for control PKM1, were stably overexpressed in RPE-1 cells and their aggregation was assessed in non-stressed cells (untreated) or cells exposed to the pH-lowering drug DMA. To ascertain comparable expression levels of all overexpressed proteins, GFP-expressing cells were FACS sorted and protein levels were confirmed by Western blotting (Extended Data Fig. 4C). As expected, acidification of the cytoplasm upon DMA treatment caused the formation of PKM2-GFP foci, while PKM1-GFP remained soluble (Fig. 4D). Interestingly, GFP-tagged PKM2^H391Y^ did not aggregate at low cytosolic pH, while PKM2^H391R^ formed constitutive aggregates independent of cytosolic pH (Fig. 4D). Together, *in vitro* and *in vivo* data demonstrate that a positive charge at position 391 in the amyloid core is both necessary and sufficient to trigger PKM2 aggregation.

### Conserved molecular principles for pH-regulated amyloid reversibility of pyruvate kinases

In summary, our findings demonstrate that cells use pH-sensing amyloid cores to regulate the formation and disassembly of functional pyruvate kinase amyloids, revealing a striking mechanism that is conserved from yeast to humans. Indeed, we show how physiological changes in cytosolic pH modify the charge of specific protonatable residues, glutamic acid in yeast and histidine in human cells, thereby regulating amyloid fibril formation and disassembly. This contrasts with the rather hydrophobic cores of aberrant, irreversible amyloids, which lack this regulatory mechanism^20^. While both PKM2 and Cdc19 contain pH-sensing residues, their sequences are surprisingly different. Thus, although the mechanism of pH-sensing has been maintained throughout evolution, the specific residues responsible for pH-sensing have been adapted, most likely to adjust to the less-pronounced cytosolic pH changes observed in stressed mammalian cells^24, 29^. In yeast, the PK amyloid core senses pH changes *via* two glutamic acids (pK_a_ 4.2 ± 0.9^38^) that become protonated and thus uncharged upon stress-induced acidification of the cytoplasm (Fig. 4E). In human cells, the PK amyloid core instead contains histidine (pK_a_ 6.6 ± 1.0^38^), which gets positively charged upon protonation. Nevertheless, in either case these protonation events result in a net charge of 0 of the amyloid core, which is likely a prerequisite for adopting a β-sheet-rich amyloid structure. Once stress is released, return to neutral cytosolic pH and the consequent deprotonation of the relevant glutamic acid and histidine residues destabilizes the amyloid cores and thereby promotes fibril disassembly. While this pH-regulation is critical, additional mechanisms cooperate to promote rapid amyloid disassembly *in vivo*. For example, disassembly of Cdc19 amyloids in yeast is triggered by binding to the allosteric regulator fructose-1,6-biphosphate (FBP), which in turn triggers a conformational change to recruit dedicated chaperones^7^. Since PKM2 but not PKM1 is allosterically activated by FBP^37^, it is likely that this mechanism also couples metabolism and PKM2 aggregation in mammalian cells. Irrespective, disassembly of Cdc19 amyloids is critical to restart energy production and thus likely also fuels the increase of cytosolic pH by providing sufficient ATP to activate specific pumps such as Pma1 (NHE1 in mammals) that secrete protons into the extracellular environment. In addition to the direct effects of cytosolic pH, amyloid reversibility may further be affected by viscosity changes that are influenced by cytosolic pH^29^.

Importantly, our results highlight a plausible role of reversible PKM2 aggregation in disease settings such as cancer. Indeed, several cancer cell types are characterized by strong upregulation of PKM2 expression, which is thought to favour cancer metabolism and contribute to the Warburg effect, while deletion of PKM2 has been shown to slow tumor growth^7^. Interestingly, it was recently reported that PKM2 activity is not necessary for cancer cell proliferation, which may rather be driven by the inactive state of PKM2, while non-proliferating tumor cells require active pyruvate kinase^30^. PKM2 amyloids are catalytically inactive, and it is thus tempting to speculate that PKM2 aggregates might play a role in cancer progression. Moreover, since the patient-derived H391Y mutation was characterized by its inability to form reversible PKM2 aggregates upon stress, it would be interesting to further explore its disease relevance.

Finally, we speculate that protonation of specific residues within amyloid cores could not only regulate reversible aggregation of pyruvate kinases, but may be a widespread mechanism to control functional amyloid formation and disassembly. Indeed, beyond pyruvate kinases, changes in cytosolic pH have been shown to influence aggregation of other functional amyloids such as peptide hormones^39^, neuropeptides^40^ and the memory-associated protein Orb2^4^. For the latter, pH-sensing was proposed to be mediated by histidine residues located in the Orb2 amyloid core^22^. Importantly, changes in intracellular pH regulate both physiological and pathological cellular processes. On the one hand, pH changes have been reported in response to cellular stresses such as starvation and heat shock both in yeast and mammalian cells^23, 28, 41, 42^, as well as during cell proliferation, cell cycle progression and differentiation^43^. On the other hand, pH changes have een associated with oth normal rain aging and Alzheimer’s disease^44^, as well as with amyotrophic lateral sclerosis (ALS)^45^. Thus, pH-sensing amyloid cores could act as pivotal and conserved effectors, directly coupling functional, reversible amyloid formation with different cellular processes and play a role in disease pathogenesis.

## Materials and Methods

### Protein purification

Protein purification was performed as previously described^1, 2^. Briefly, *E. coli* cells (Rosetta) were transformed with plasmids expressing either wild-type or mutant Cdc19, PKM1, or PKM2. Cells were grown at 37 °C in LB media (1 % peptone, 0.5 % yeast extract, 0.5 % NaCl) containing 30 µg/ml chloramphenicol and 100 µg/ml carbenicillin until reaching OD_600_ 0.6. Then, IPTG was added to a final concentration of 0.1 mM to induce protein expression. Cells were grown at 16 °C for 12 h, harvested by centrifugation, resuspended in cold purification uffer (100 m Tris-HCl pH 7.4, 200 m NaCl, 1 m gCl_2_, 10 % glycerol, 1 m phenylmethylsulfonyl fluoride (P SF), 1 m DTT) supplemented with protease inhi itor ta lets (Roche, 11697498001) and 75 U/ml of Pierce universal nuclease (Thermo Fisher Scientific, 88700), and lysed by freezer milling (SPEX SamplePrep 6870 Freezer/Mill; five cycles of 2 min cooling and 2 min grinding at setting 15 CPS). For PK 1, wild-type and mutant PKM2 purifications, the purification buffer contained 20 % glycerol. After clearing the lysates by centrifugation (4 °C, 30 min, 48000 g), the supernatant was loaded on a Strep-Tactin Superflow Plus column (Qiagen) at 4 °C following the manufacturer’s instructions. Proteins were eluted using purification buffer supplemented with 2.5 mM D-desthiobiotin, their purity was checked by SDS-PAGE and Coomassie blue staining, and pure aliquots were stored at -80 °C.

### Prediction of amyloidogenic regions, peptide selection and fibrils preparation

The amino acid sequence of Cdc19 (Saccharomyces Genome Database SGD identifier: S000000036, https://www.yeastgenome.org/locus/S000000036) was submitted to the amyloid- predicting consensus tool AmylPred2.0 (http://aias.biol.uoa.gr/AMYLPRED2/3), and the structure-based prediction tool ZipperDB (http://services.mbi.ucla.edu/zipperdb/4). ZipperDB calculates the fibrillation propensities for every possible hexapeptide in the protein sequence of interest, while AmylPred2.0 combines the prediction of 11 different methods developed to identify regions likely to form amyloid fibrils. Based on these predictions, we selected 20 hexapeptides, distributed over the whole Cdc19 sequence, which included all regions predicted to be particularly amyloidogenic and four non-amyloidogenic negative controls. The amyloid core of Cdc19 corresponding to amino acids 376-392 was defined as the region that was experimentally validated to readily form amyloids (based on the above-mentioned hexapeptide screening) and predicted to be an aggregation-prone LCR using the SEG program (http://mendel.imp.ac.at/METHODS/seg.server.html5). The PKM2 amyloid core was defined as the region predicted to be an aggregation-prone LCR by the SEG program^5^, and correspond to amino acids 372-402 in PKM2. Also a shorter PKM2 amyloid core (containing only one histidine instead of two) was analysed, and corresponds to amino acids 382-402 in PKM2. All above-mentioned peptides were ordered in lyophilized form from GL Biochem, dissolved in DMSO with 10% formic acid to a concentration of 10 mg/ml, and stored at -20 °C until use. To prepare fibrils, the hexapeptide stocks were diluted to 2 mg/ml in de-ionized H_2_O and incubated over night at 30 °C. To prepare fibrils of the Cdc19 and PKM2 amyloid cores, the peptides were incubated at the indicated pH (see legends) in PBS 1x or Tris-HCl buffer (100 mM Tris-HCl, 200 m NaCl, 1 m gCl_2_), for two days at 30 °C at a final concentration of 2 mg/ml. Incubation in PBS or Tris yielded equal fibril morphologies in TEM. For CD measurements, PBS 1x buffer was used.

### Transmission electron microscopy (TEM)

TEM images were acquired on a FEI Morgagni 268 electron microscope at 100 kV using a CCD 1376 x 1032 pixel camera at different magnifications. Peptide fibrils were obtained as described a ove and 5 μl of the sample was spotted on non-glow discharged carbon film 300 mesh copper grids (CF300-CU from Electron Microscopy Sciences) and incubated for 1 min. Purified full-length Cdc19, PKM1 and PKM2 aliquots were thawed on ice and cleared by centrifugation (4 °C, 10 min, 21000 g). Samples were then diluted to 0.3 mg/ml in purification buffer with a final pH of 6, and 5 μl of the sample was spotted on non-glow discharged grids and incubated for 10 min at 4 °C or 42 °C. For both peptides and full-length proteins, the excess sample was manually blotted with Whatman filter paper, and the grid was washed twice with the same buffer in which the proteins or peptides were dissolved. The grid was then negatively stained with two drops of 2 % uranyl acetate and air dried.

### Atomic force microscopy (AFM)

Cdc19 amyloid core fibril solution was obtained as described above and diluted to 0.05 mg/ml. The freshly cleaved mica was functionalized with 1 % APTES (10 µl) for 1.5 min, rinsed with Milli-Q water and dried by compressed gas. Then, an aliquot (10 µl) of diluted fibril solution was deposited on the functionalized mica for 2 min, rinsed with Milli-Q water and dried by a gentle flow of compressed gas. AFM measurements were carried out using a Bruker multimode 8 AFM (Bruker, U.S.A.) with an acoustic hood to minimize vibrational noise. AFM imaging was operated in soft tapping mode under the ambient condition, using a commercial silicon nitride cantilever (Bruker, U.S.A.) at a vibration frequency of 70 kHz. AFM images were flattened using Nanoscope 8.1 software (Bruker, U.S.A.), and no further image processing was applied.

### Cryo-EM sample preparation and data processing

The Cdc19 amyloid core peptide stock was diluted to 0.4 mg/ml in H_2_O and incubated for 4 days at 25 °C while shaking at 600 rpm. 4 µl of sample was then applied onto glow discharged Quantifoil grids and plunge frozen using a Leica Plunge Freezer system at 80 % humidity and 20 °C in a liquid ethane-propane mixture. Micrographs were acquired on a Titan Krios microscope (Thermo Fisher Scientific) operated at 300 kV with a Gatan K2 Summit direct electron detector in counting mode using a slit width of 20 eV on a GIF-Quantum energy filter. 3,356 movies were collected with a calibrated pixel size of 0.82 Å. Each micrograph was dose-fractionated to 50 frames with a total dose of approximately 54 e- /Å2. The collected movies were aligned with MotionCor2^6^, followed by the CTF determination in GCTF 1.06^7^. All subsequent image processing was performed in Relion 3.1^8^. Manual particle picking from non-overlapping narrow fibrils (Fig. 1D^A^) resulted in extraction of 263,819 segments with an interbox distance of 14.3 Å and a box size of 352 pixels. The particles were subjected to the reference-free 2D-classification.

One of the classes with characteristic pattern of the strands in β-sheets was used to estimate the rise of 4.77 Å (by measuring the distance to the peak in the average image of the power spectra of each class-average member). The estimation of the cross-over distance was hindered by the sample heterogeneity and lack of non-overlapping straight fibrils, which, in turn, hampered the 3D-analysis.

### Solid-state nuclear magnetic resonance

An Applied Biosystems 433 A automated batch peptide synthesizer was used to synthesize the Cdc19 amyloid core peptide (^376^TSTTETVAASAVAAVFE^392^) with ^13^C/^15^N labelled Glu380. The synthesis was started from commercial available Fmoc-Glu-Wang resin. The cleavage from the resin was realized with TFA / TIS / H_2_O 95:2.5:2.5 (v/v). The TFA was vaporized and the crude peptide was washed with diethyl-ether. The peptide was dried under vacuum and dissolved in DMSO and formic acid (10 %). To fibrillize the peptide, the sample was diluted with H_2_O, the pH was adjusted to 5.8, and the fibrils were formed over seven days at room temperature. Subsequently, the fibrils were centrifuged and washed (pH 5.8). Solid-state NMR rotors were filled overnight in an ultracentrifuge (16 h at 4 °C at 210’000 g) using home-build rotor-filling tools^9^. ^13^C solid-state NMR spectra of the only ^13^C-labelled E380 Core^WT^ peptide were recorded on two samples with pH values of 5.8 and 6.2, respectively. Experiments were performed at 20.0 T in a 3.2 mm triple-resonance probe using a magic-angle spinning frequency of 17.0 kHz. The spectra were recorded using adiabatic ^1^H,^13^C cross-polarization with radio frequency fields of 60 kHz (^1^H) and 45/38 kHz (^13^C) for the two samples. The CP contact time was set to 500 μs. 90 kHz ^1^H SPINAL-64 decoupling was applied during detection. The repetition time was set to 2.5 s with an acquisition time of 15 ms. For the pH 4.2 sample, 580 scans (24 min total measurement time) and for the pH 6.2 sample 2160 scans (90 min measurement time) were collected. The spectra were recorded at 278 K. Spectra were referenced to 4,4-dimethyl-4- silapentane-1-sulfonic acid (DSS) using the methylene resonance of solid adamantane as an external standard^10^. Processing of NMR spectra was performed with TOPSPIN (version 3.5, Bruker Biospin).

### Circular dichroism (CD)

CD spectra were recorded on a J-815 CD Spectrometer (Jasco) using a quartz cuvette with 1 mm path length (HellmaAnalytics, Art. No. 110-1-40) at 25 °C. Fibrils were prepared as described above, collected by centrifugation (10 min, 21000 g), and washed twice with PBS 1x, before CD measurements.

### Kinetics measurements of amyloid fibrils re-solubilization

To measure the kinetics of fibrils re-solubilization on short time scales, we designed and fabricated a microfluidic chip containing three inlets: a first inlet to inject ThT-stained, pre- formed fibrils as well as two further inlets for buffer solutions at high (pH 7.4) and low (pH 5.8) pH values. To trap the pre-formed aggregates, a wide channel containing C-shaped traps has been additionally included into the design. The Master wafers and PDMS-based microfluidic devices have been fabricated as described using standard soft lithography^11^. To operate the chip, the three aforementioned solutions were filled into 500 µl (buffers) and 100 µl (pre-formed aggregates) Hamilton glass syringes and the flow rate was controlled by Nemesys syringe pumps (Cetoni, Germany). At first, the fibrils pre-formed in a buffer at pH 5.8 were injected into the chip and trapped using the integrated C-traps. Subsequently, the chip was flushed with the same buffer at pH 5.8 to remove non-trapped, residual fibrils from the channels. To then dissolve the trapped aggregates, the high-pH buffer was flushed at flow rates of 5 µl/min. Aggregate re-solubilization was monitored by recording ThT fluorescence (excitation: 450 nm, emission: 490 nm) over time of at least five trapped aggregates simultaneously using a Nikon TI Eclipse Microscope equipped with an Andor Zyla camera and an Omicron LED Hub laser source. To extract the characteristic time τ of aggregate dissolution, the intensities of single aggregates trapped in separate traps were extracted over time by using an in-house written Matlab code. Briefly, to distinguish the fluorescence signal of the aggregates from the background, the mean intensity of each image was defined as threshold. The mean background intensity was then subtracted from each image and the corrected signal was averaged for each image, normalized and plotted over time. The resulting normalized signal I_norm_ was fitted to equation (1) to obtain the characteristic time τ.

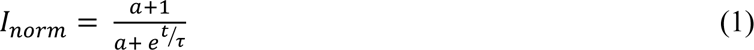

### ThioflavinT (ThT) and Congo Red (CR) staining

Thioflavin T (ThT, Sigma-Aldrich, T3516) or Congo Red (CR, Sigma-Aldrich, 75768) were dissolved in water to a final concentration of 2.5 mM or 1 mM, respectively, and filtered (0.2 µm filter, Millipore). Either ThT or CR were then mixed (1:10 dilution) with fibrillized peptide samples in a 384-well plate (Corning Life Sciences). Full-length Cdc19 in purification buffer was thawed on ice, cleared by centrifugation (10 min, 4 °C, 21000 g), and adjusted to a concentration of 0.3 mg/ml and pH as indicated in the figure legend prior to ThT or CR addition. ThT and CR signals were measured in a CLARIOstar plate reader (BMG Labtech), with 450 nm excitation and 490 nm emission, and excitation at 560 nm and emission at 614 nm, respectively.

### Semi-denaturing detergent agarose gel electrophoresis (SDD-AGE)

The indicated yeast strains were grown in 5 ml at 30 °C, harvested after the indicated growth period, cells were washed once with water and resuspended in ∼300 µl ice-cold lysis buffer (50 mM Tris pH 7.5, 150 mM NaCl, 1 % (vol/vol) TritonX-100, 2.5 mM EDTA, 0.33 mM PMSF, protease inhibitor tablet (Roche, 11697498001), 6.7 mM NEM). The mixture was added to ice- cold glass beads and the cells were lysed using mechanical disruption (6 m/s for three times 20 s with 5 min pause). After centrifugation, the supernatant samples were adjusted for equal protein concentrations and mixed 4:1 with 4 × Sample buffer (40 mM Tris acetic acid, 2 mM EDTA, 20 % glycerol, 4 % SDS, bromophenol blue). Samples were incubated for 10 min at room temperature and loaded onto a 1.5% agarose gel containing 0.1% SDS in 1 × TAE/0.1% SDS running buffer. The gel was run at low voltage or in the cold. Proteins were detected by immunoblotting with a GFP-specific antibody.

### Size-exclusion chromatography (SEC)

Purified wild-type or mutant Cdc19 were thawed on ice, and 0.1 mg protein were loaded on a Superdex 200 10/300 GL size-exclusion column (GE Healthcare) connected to an ÄKTA pure (GE Healthcare) at 4 °C. The column was previously equili rated in uffer (100 m Tris-HCl pH 7.4, 200 m NaCl, 1 m gCl_2_, 10 % glycerol) and run according to manufacturer’s instructions. Protein elution was followed by measuring UV absorbance (280 and 215 nm, a.u.).

### Pyruvate activity assay

Pyruvate kinase activity was measured as previously described^2^. Briefly, the pyruvate kinase reaction was coupled to the lactate dehydrogenase reaction and assayed by spectrophotometrically measuring the conversion of NADH to NAD+ at 340 nm. Purified PKM1 or PKM2 was thawed on ice, cleared by centrifugation (4 °C, 10 min, 21000 g), diluted to 0.2 mg/ml in purification buffer, and either kept on ice (soluble) or heat shocked for 3 h at 45 °C. Soluble and aggregated protein was diluted in activity buffer (50 mM imidazole pH 7, 100 mM KCl, 25 mM MgCl_2_, 10 mM ADP, 0.3 mM NADH, 10 U/ml LDH) to a final protein concentration of 2 μg/ml. Reactions were started y adding PEP (final concentration 2 m), and decrease in absorbance at 340 nm was monitored over time.

### Yeast cell growth and fluorescence microscopy

Yeast strains used in this study are listed in Supplementary Table S3. Cells were grown in synthetic SD media (2 % glucose, 0.5 % NH_4_-sulfate, 0.17 % yeast nitrogen base, and amino acids) at 30 °C. Growth was observed by spotting cells in serial dilutions on SD agar plates, and imaging the plates after 3 days at 30°C. Fluorescence microscopy was performed using a Nikon Eclipse Ti-E microscope with MicroManager software. For time-lapse experiments, exponentially growing yeast cells (OD_600_ 0.4-0.6) were loaded in commercial microfluidic chips (CellASIC ONIX2, Merck Millipore) as previously described ^2^, and images were recorded every 10 min. For glucose starvation experiments, starvation media (i.e. synthetic SD media without glucose: 0.5 % NH_4_-sulfate, 0.17 % yeast nitrogen base, and amino acids) was supplemented with Alexa Fluor 647-Dextran (10,000 MW, Invitrogen) to control successful switch of media.

### Molecular biology

Plasmids used in this study are listed in Supplementary Table S4. DNA mutations were introduced by site-directed mutagenesis using standard molecular biology protocols. Sequences of PKM1 and PKM2 were retrieved from p413TEF-PKM1 and p413TEF-PKM2 plasmids (Addgene, 34607 and 34608), and Gibson assembly was performed to clone PKM1, PKM2 and PKM2 mutants into pLenti-CMV-MCS-GFP-SV-puro (Addgene, 73582).

### Protein levels quantification

To prepare total protein extracts, exponentially growing yeast cells expressing wild-type or mutant Cdc19-GFP were treated with 10 % trichloroacetic acid (TCA) and incubated on ice for at least 10 min. Cells were harvested by centrifugation and washed twice with ice-cold acetone. Then, acetone was removed, pellets were resuspended in 8 M urea sample buffer and boiled for 10 min at 70 °C. Samples were analysed by western blotting using a α-GFP antibody (Roche, 11 814 460 001), and a α-Pgk1 antibody (Invitrogen, 459250) as control.

### Human cell culture and RNAi-depletion

RPE-1 cells were maintained in DMEM media supplemented with FCS (10 % final concentration) and Penicillin Streptavidin-Glutamine (PSG; Gibco, 1 % final concentration). For starvation, cells were washed once with RPMI medium (Gibco, w/o glucose, FCS and PSG) and incubated in the same medium for 24 hours. Cells were stimulated with RPMI medium containing FCS (10 % final) and Glucose (5 mg/ml) as indicated. Where indicated, cycloheximide (CHX, Sigma-Aldrich, 01810) or dimethyl amiloride (DMA, Sigma-Aldrich, A4562) were added to the medium at a concentration of 1 μ or 100 μ, respectively. RNAi to deplete NHE1 or PKM2 was performed using RNAiMAX transfection reagent (Thermo Fisher Scientific, 13778100) following the manufacturer’s instructions. siRNA sequences used in this work were verified as described in ^12^ and are: si_155 (5’-GCC AUA AUC GUC CUC ACC A), si_156 (5’-CC AUA AUC GUC CUC ACC AA), si_27 (5’-AGG CAG AGG CUG CCA UCU A).

### Lentivirus generation and transduction

Lentivirus generation was conducted following a standard protocol. Briefly, HEK293T cells were co-transfected with a plasmid encoding the lentiviral envelope (pMD2.G), a second-generation lentiviral packaging plasmid (psPAX2), and the target plasmid using Lipofectamine2000. 6-8 h post-transfection, the media was changed, and the lentivirus was harvested by filtering the supernatant with a 45 µm filter. For transduction, the lentivirus was added to the cell line of interest at a 1:100 dilution. Then, cell lines were passaged 5 times and sorted for equal GFP-levels using fluorescence-activated cell sorting FACS.

### Immunofluorescence

Immunofluorescence was performed essentially as previously described^13^. In brief, cells were grown on glass coverslips, washed with PBS 1x, fixed in 4 % paraformaldehyde for 20 min at room temperature. Permeabilization was performed adding 0.1 % Triton-X100 in PBS 1x for 10 min at room temperature, followed by 3x washes in 0.01 % Triton-X100 in PBS 1x (washing buffer). The cells were then incubated with 3 % BSA in washing buffer (blocking buffer) for 20 min to 1 hour at room temperature. Primary α-PKM1 antibodies (Cell Signaling Technologies, (D30G6) XP® Rabbit mAb #7067), and α-PKM2 antibodies (Cell Signaling Technologies, (D78A4) XP® Rabbit mAb #4053) were diluted (1:3000) in blocking buffer and incubated for 1 hour at room temperature. After 3x washes with washing buffer, the cells were incubated with secondary antibody (Alexa Fluor-conjugated anti-rabbit/or anti-mouse IgG (Thermo Fischer Scientific) diluted in blocking buffer for 1 hour at room temperature. Nuclei staining was performed by applying 0.2 µg/ml DAPI (Sigma-Aldrich, D9542) in washing buffer for 10 min at room temperature. After 3x washes in washing buffer, the coverslips were mounted onto microscopy slides using Immu-Mount (Thermo Fischer Scientific). The images were captured on an inverted Ti-Eclipse microscope (Nikon) with either 40x oil objectives or 60x objectives and MicroManager, V 1.4. Images were analyzed using FIJI (ImageJ V 2.0.0).

### Pelleting assays

Purified proteins were thawed on ice and cleared by centrifugation (4 °C, 10 min, 21000 g), and diluted to a final protein concentration of 0.5 mg/ml in 100 m Tris/HCl pH 7.4, 200 m NaCl, 1 m gCl_2_, 10% glycerol, 1 m DTT, 1 m P SF. Proteins were kept on ice or heat shocked at 42 °C for 10 or 20 min, as indicated in the legends. Aggregates were pelleted by centrifugation (4 °C, 10 min, 21000 g), and separated from the supernatant containing soluble protein. Aggregation was quantified by loading pellet and supernatant on a SDS-PAGE gel. RPE-1 cells were seeded on 15 cm plates and allowed to attach overnight. Then, two plates were washed with 1x PBS or RPMI media without supplements, and cultured in RPMI without supplements for 24 h (starved sample), while the rest was left untreated. Subsequently, cells were washed with PBS 1x prior to collection by centrifugation (500 g, 5 min, room temperature). Pellets were resuspended in lysis buffer, and lysed on ice for 30 min. Resulting lysates were cleared y centrifugation (10 min, 10’000 rpm, 4 °C), and protein concentration was adjusted in untreated and starved samples by Bradford measurements. Cleared lysates were centrifuged (20 min, 14’000 rpm, 4 °C), supernatant was separated from pellet, and analysed y Western blotting.

### Antibodies and reagents

For western blotting, the following antibodies were used (all at 1:3000 dilution): α-GFP (Roche, 11 814 460 001), α-Pgk1 (Invitrogen, 459250), α-Vinculin (Sigma-Aldrich, V9131), HRP-coupled secondary antibody (Biorad, 170-6516), α-PKM1 (Cell Signaling Technologies, (D30G6) XP® Rabbit mAb #7067), α-PKM2 (Cell Signaling Technologies, (D78A4) XP® Rabbit mAb #4053). For immunofluorescence, the above-mentioned α-PK 1 and α-PKM2 antibodies were used.

### Statistics and Reproducibility

All data are representative results from at least three independent experiments, unless differently specified in the figure legends. GraphPad Prism was used to analyse and plot the results, and whenever possible mean ± S.E.M. and individual data points of individual experiments are shown. No outlier tests were performed and no data were excluded from the analyses. Statistical tests used and *P* values are indicated in the respective figure legends.

### Data availability

Yeast strains, human cell lines, plasmids and reagents, as well as detailed experimental procedures and additional data supporting the findings of this study are available from the corresponding authors upon request.

### Code availability

No custom code was used in this work.

## Acknowledgments

We thank Lorenzo Garbani Marcantini for help with data analysis, the Scientific Center for Optical and Electron Microscopy (ScopeM) of ETH and in particular Miroslav Peterek for microscopy support, and Beat H. Meier for providing NMR measurement time. We are grateful to Matthew Vander Heiden, Markus Stoffel, Reinhard Dechant, Alicia Smith and members of the Peter lab for discussions and comments on the manuscript. This work was supported by the Swiss National Science Foundation, the Synapsis Foundation, the Human Frontier Science Program and ETH Zürich.

## Author contributions

Conceptualization: GC and MP; Formal analysis: GC; Funding acquisition: GC and MP; Investigation: GC, VK, LK, AA, PAf, ML, CE, JZ, YC, DP, SK, TW, RC; Software: PAf; Supervision: DB, RM, PAr, RR, MP; Visualization: GC; Writing - original draft: GC; Writing – review and editing: GC, MP, with inputs from all co-authors.

## Competing interests

The authors declare no competing interests.

## Supplementary information

Lists of plasmids and strains used in this study are reported in Supplementary Table S3 and Supplementary Table S4.

## Corresponding authors

Correspondence and requests for materials should be addressed to Gea Cereghetti or Matthias Peter.

## Extended data figures and tables

**Extended Data Fig. 1.**
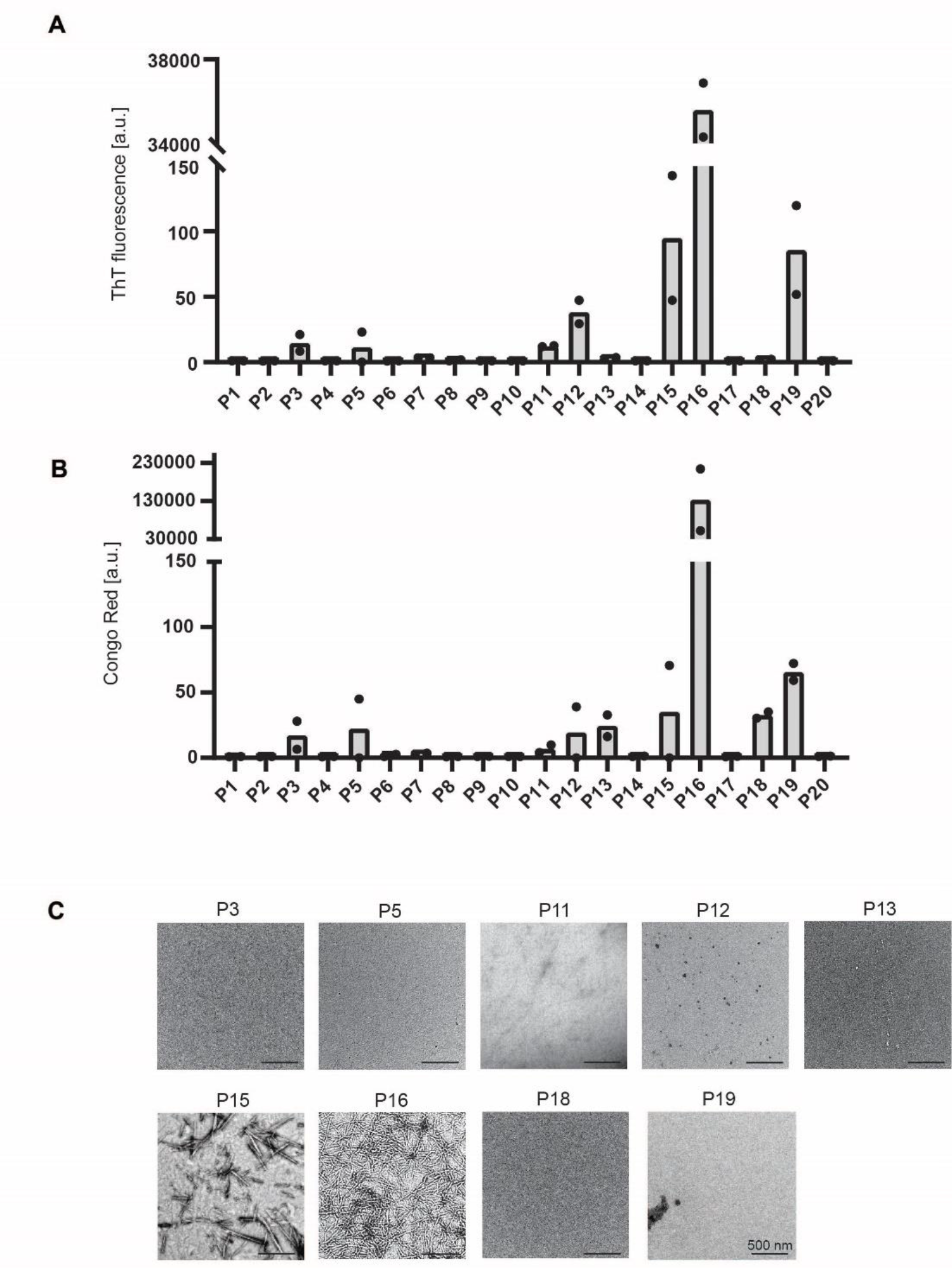
Peptide-based screening to identify the amyloid core of Cdc19. (A)- (B) Identification of amyloidogenic regions within Cdc19. Based on the amyloid prediction tools ZipperDB^10^ and AmylPred2.0^9^, 16 hexapeptides corresponding to the regions with highest amyloidogenicity (+ 4 negative controls) were selected and screened for their ability to form amyloids by Thioflavin T (A) and Congo Red (B) staining (n = 2 independent experiments). (C) Validation of screen hits by negative staining TEM. ThT- and/or CR-positive peptides were visualized by negative staining TEM to confirm their ability to form fibrillary amyloid-like structures (n= 3). Scale bar: 500 nm.

**Extended Data Fig. 2.**
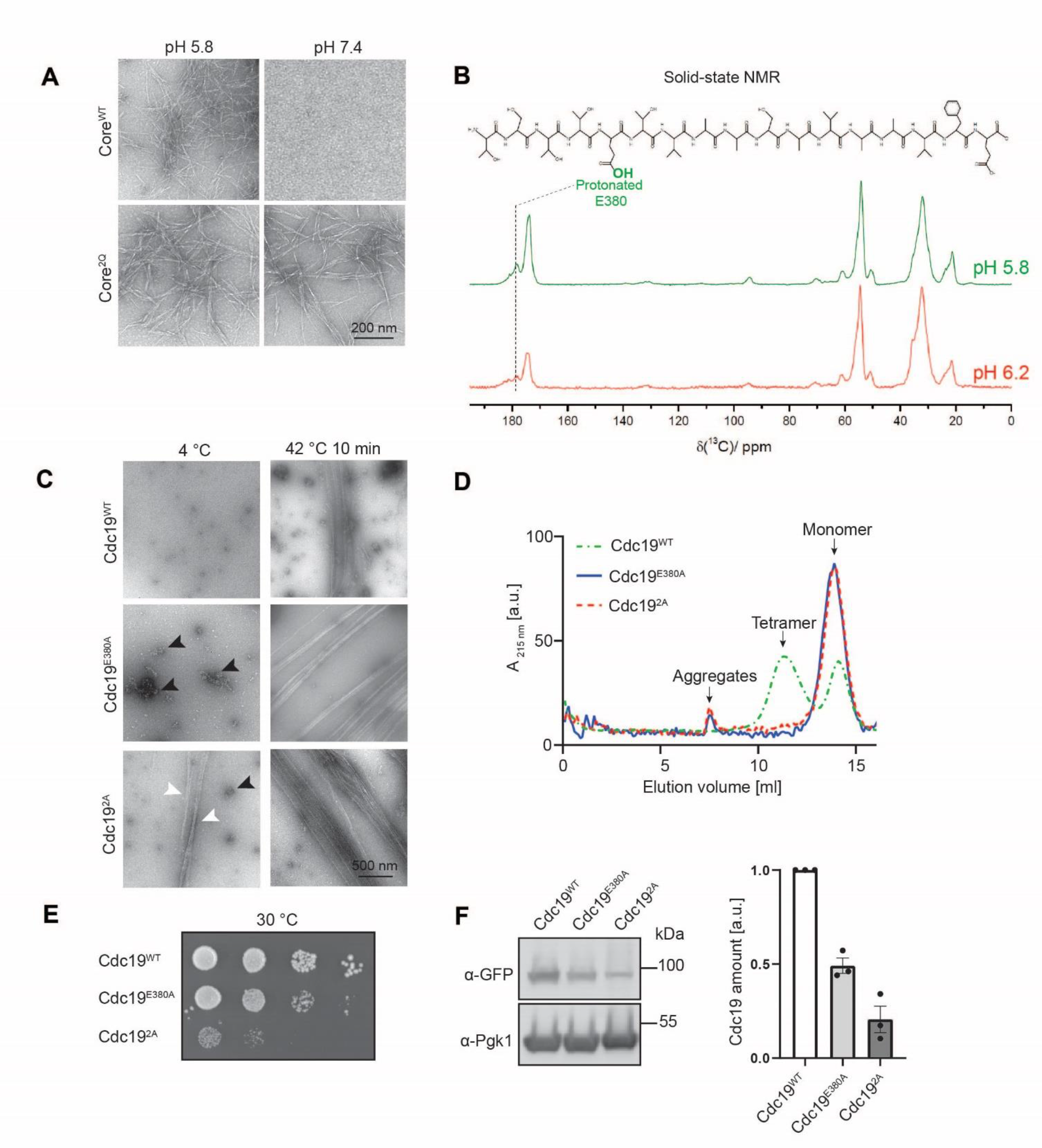
E380 and E392 in the Cdc19 amyloid core sense pH and regulate pH-dependent amyloid formation. (A) Cdc19 amyloid core peptides forms amyloid fibrils only at physiologically low pH. Cdc19 wild-type (Core^WT^) or mutant (Core^2Q^, mutations: E380Q, E392Q) amyloid core peptides were incubated at the indicated pH for two days and imaged by negative staining TEM. Note that Core^WT^ forms fibrillar aggregates at physiologically low pH (corresponding to the intracellular pH of stressed cells), while it remains soluble at neutral pH (corresponding to the intracellular pH of growing cells). Core^2Q^ instead is pH-insensitive and forms fibrils under both conditions. n = 3. Scale bar: 200 nm. (B) ^13^C-solid-state NMR spectra of the Core^WT^ peptide at pH 5.8 or pH 6.2. The pH-sensing glutamic acid E380 is protonated in the amyloid Cdc19 core at pH 5.8 and gets partially deprotonated upon pH increase to 6.2 as judged by chemical-shift changes of the carboxyl carbon atom. The chemical structure of the peptide is shown, and the relevant peak of the carboxyl carbon sensitive to protonation of the ^13^C- labelled E380 residue is highlighted. (C) Purified full-length Cdc19 mutants (Cdc19^E380A^ and Cdc19^2A^) rapidly form amyloid fibrils independently of pH. Cdc19^E380A^ and Cdc19^2A^ were recombinantly expressed and purified from *E. coli*. The yield was very low compared to wild-type controls as most of the protein aggregated during purification. In contrast to wild-type controls, the small amounts that could be purified rapidly formed large oligomers (black arrow heads) or fibrils (white arrow heads) already at 4 °C, pH 7.5 (n = 3). Scale bar: 500 nm. (D) Purified full-length pH-insensitive Cdc19 mutants (Cdc19^E380A^ and Cdc19^2A^) are aggregation-prone independently of pH. Size-exclusion chromatography (SEC) of freshly purified Cdc19^WT^, Cdc19^E380A^ and Cdc19^2A^ indicates that Cdc19^WT^ is present as a mixture of stable tetramers and monomers, while Cdc19^E380A^ and Cdc19^2A^ are exclusively present in the aggregation-prone monomeric form, and tend to form large aggregates even at pH 7.5 and 4 °C. n = 3. (E) Cdc19 mutants with impaired pH-sensing exhibit growth defects. Yeast cells expressing wild-type (Cdc19^WT^) or the Cdc19^E380A^ or Cdc19^2A^ mutants were grown at 30 °C. Serial dilutions were then spotted on agar plates and grown at 30 °C for 3 days to observe their growth rate (n = 3). (F) Quantification of Cdc19 protein levels. Cells expressing GFP-tagged wild-type (Cdc19^WT^) or Cdc19^E380A^ or Cdc19^2A^ mutants were lysed and immunoblotted with antibodies against GFP (top panel) or for control Pgk1 (bottom panel). Mean Cdc19- GFP levels normalized with Pgk1 are shown (n = 3).

**Extended Data Fig. 3.**
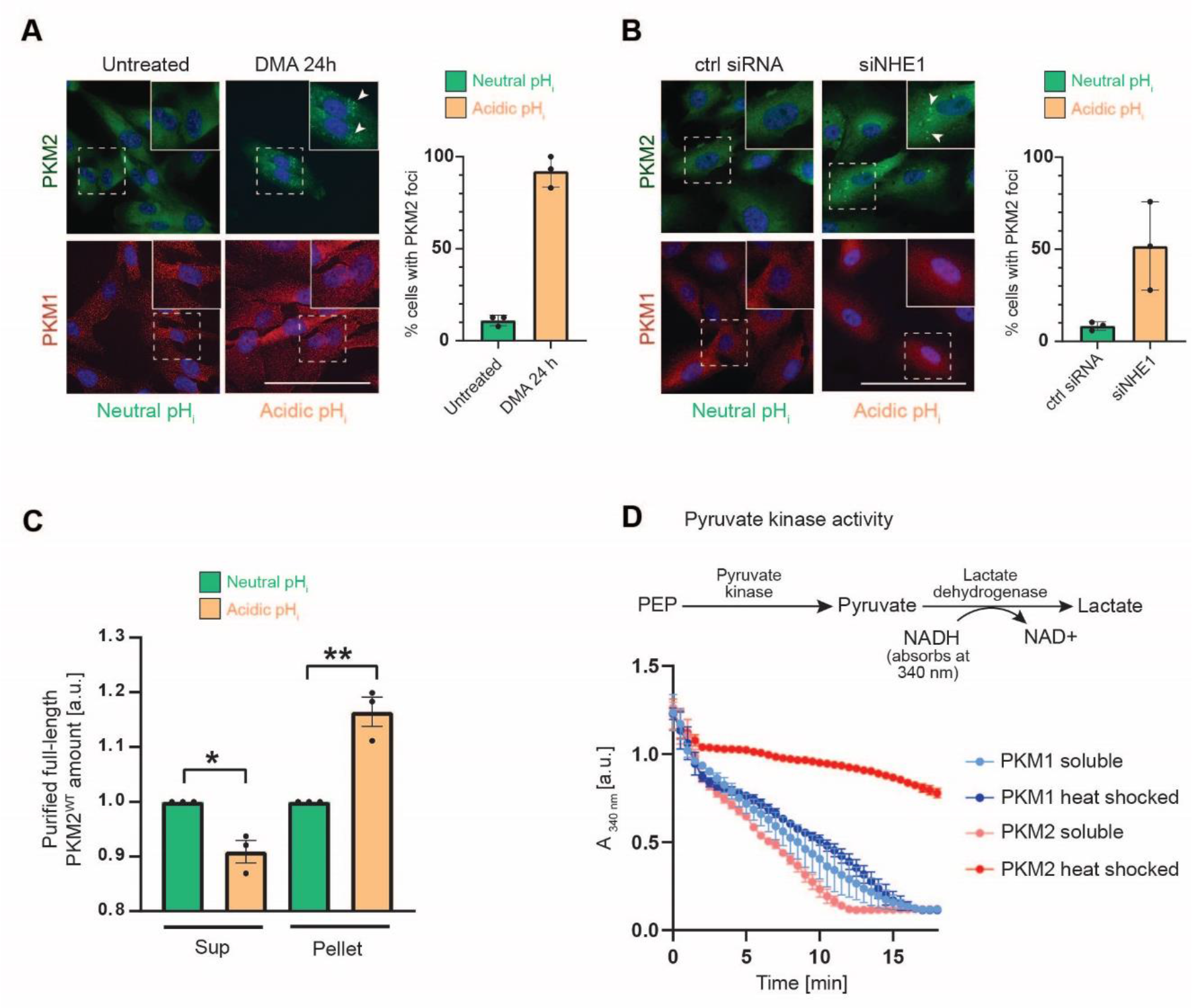
Decreasing cytosolic pH triggers PKM2 aggregation, which inactivates the protein. (A) Formation of PKM2 but not PKM1 foci upon DMA-induced decrease of cytosolic pH. RPE-1 cells were left untreated or treated with the pH-lowering drug D A (100 μ) for 24 h. The localization of PKM2 and PKM1 was analysed by immunofluorescence and the percentage (%) of cells with cytoplasmic PKM2 foci was quantified as mean ± SEM (n = 3). At least 50 cells were analysed for each condition. Scale ar: 50 μm. (B) Artificially lowering cytosolic pH by siRNA-depleting NHE1 triggers aggregation of PKM2 but not PKM1. RPE-1 cells were subjected to siRNA against NHE1 or control siRNA. PKM2 and PKM1 localization was analysed by immunofluorescence and the percentage (%) of cells with cytoplasmic PKM2 foci was quantified as mean ± SEM of three independent experiments. At least 50 cells were analysed for each condition. Scale ar: 50 μm. (C) Lowering pH causes aggregation of purified full-length PKM2. Full-length PKM2^WT^ was purified at pH 7.4. Then pH was either kept constant or lowered to pH around 6 by adding HCl, and samples were incubated overnight at 4 °C. Soluble protein (Sup) was separated from aggregates (Pellet) by centrifugation, analysed by SDS-PAGE and quantified after Coomassie blue staining. The graph shows normalized PKM2 amounts as mean ± SEM (n = 3, two-tailed Student’s t-test, *\*P* = 0.0113, *\*\*P* = 0.0036). (D) Soluble PKM1 and PKM2 are catalytically active, while aggregated PKM2 is inactive. Pyruvate kinase activity was measured using a lactate dehydrogenase-coupled activity assay before (soluble) or after (heat shocked) 2 h heat shock at 50 °C. Briefly, active pyruvate kinase converts phosphoenolpyruvate (PEP) into pyruvate, which in turn is reduced to lactate by lactate dehydrogenase. The concomitant conversion of NADH to NAD^+^ is assayed as decrease in absorbance at 340 nm over time, and is used as a measure of pyruvate kinase activity^7^. Mean ± SEM (n = 3 independent experiments) is shown.

**Extended Data Fig. 4.**
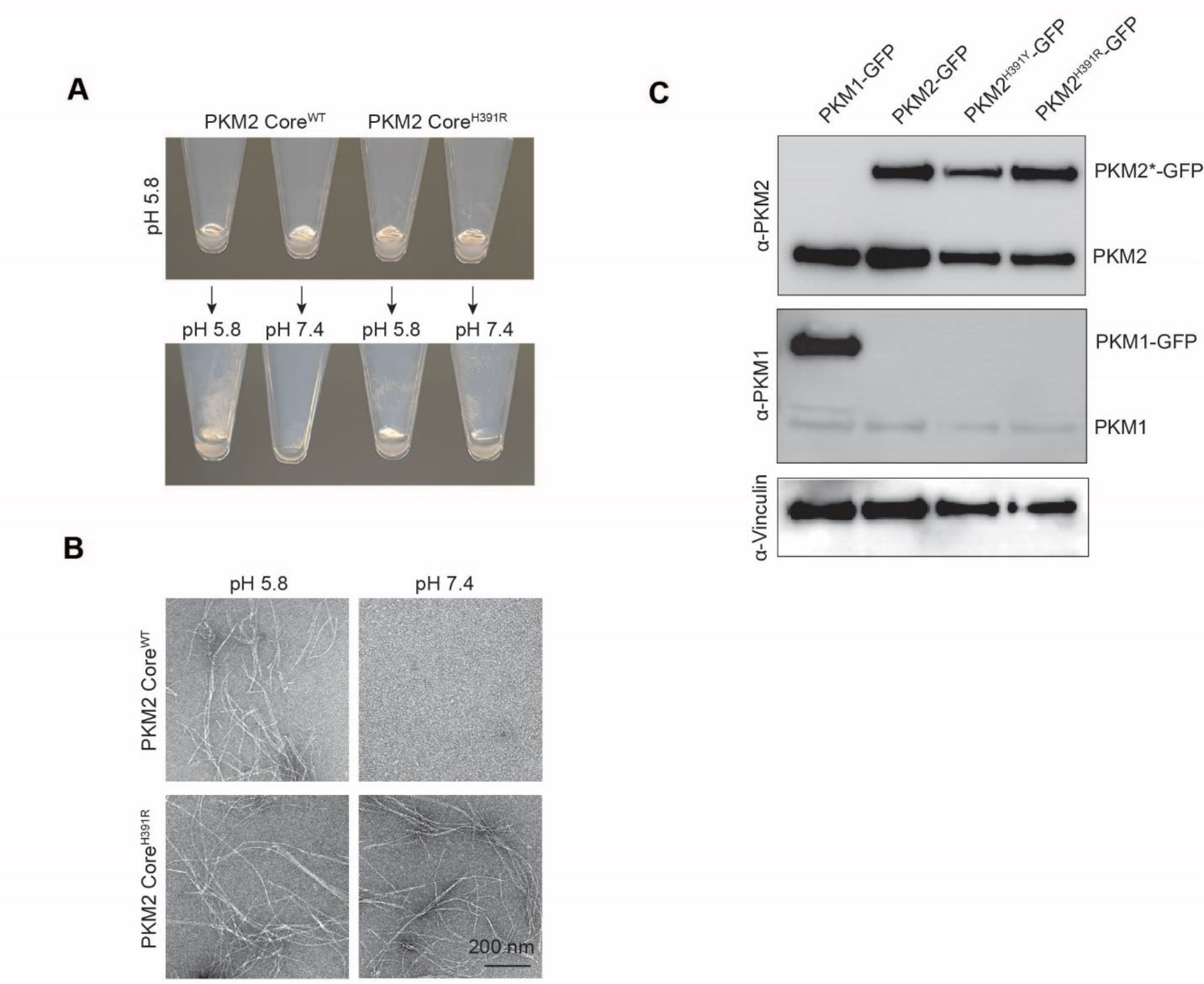
Formation and disassembly of PKM2 amyloids is regulated *via* protonation of histidine 391 (H391) in its amyloid core. (A) - (B) H391 in the amyloid core of PKM2 regulates pH-dependent aggregation. PKM2 wild-type (Core^WT^) or the Core^H391R^ mutant peptides (amino acids 382-402) were incubated at pH 5.8 overnight. Then, pH was either kept constant, or increased to pH 7.4. Samples were centrifuged and the resulting pellets photographed (A) (n = 3) or visualized by negative staining TEM (B). Note that amyloid fibrils of Core^WT^ re-solubilize at high pH (no fibrils visible by TEM), while the pH-insensitive mutant Core^H391R^ presents fibrils regardless of pH (n = 3). Scale bar: 200 nm. (C) Expression analysis of endogenous and ectopically expressed GFP-tagged wild-type and mutant PKM proteins in the indicated cell lines. GFP-tagged wild-type PKM1, PKM2 or the indicated PKM2 mutants (PKM2^H391R^ or PKM2^H391Y^) were overexpressed in RPE-1 cells. Cells were lysed and immunoblotted with antibodies against PKM2 (top panel), PKM1 (middle panel) or for control Vinculin (bottom panel). Note that expression levels of endogenous and GFP-tagged PKM2 are comparable, while GFP-tagged PKM1 is strongly overexpressed compared to endogenous PKM1.

**Extended Data Table 1.** Structural features of the Cdc19 amyloid core, compared with different pathological amyloids (i.e. α-syn^47^ and Aβ^15^). Data describing the characteristics of the Cdc19 amyloid core are shown as mean ± S.E., “n” is indicated.

| **Fibril characteristics** | **Cdc19**<br>**(376-392)** | **α-syn (1-121)**<br>**(2018)** | **Aβ(1-42)**<br>**(2015)** | **Aβ(1-42)**<br>**(2017)** |
| --- | --- | --- | --- | --- |
| **Fibril diameter [Å]** | 23 ± 4<br>(n = 35) | 100 | 100 | 70 |
| **Helical rise [Å]** | 4.77 | 2.45 | - | 4.67(C2)/2.34 |
| **Spacing between subunits [Å]** | 4.77 | 4.9 | 4.7 assumed | 4.67 |
| **Helical twist [°]** | Left-<br>handed | 179.5 (azimuthal) | -0.769 per<br>subunit | -179.3 (azimuthal) |
| **Crossover distance [Å]** | Variable:<br>1010 ± 180<br>(n = 35) | - | 1100 | - |
| **Staggered protofibrils** | yes | yes | Yes and not<br>found | yes |

**Extended Data Table 2.** Sequence composition of Cdc19 and PKM2 amyloid cores and comparison with other functional or pathological amyloid cores^20^.

|  | **Gly %** | **Asn + Gln %** | **Tyr + Ser + Thr %** | **Val + Ala + Ile + Leu %** |
| --- | --- | --- | --- | --- |
| **Cdc19**<br>(functional) | 0.0 | 0.0 | 35.0 | 47.1 |
| **PKM2** | 0.0 | 6.5 | 3.2 | 48.4 |
| **HnRNPA2-LCD**<br>(functional) | 35.1 | 22.8 | 22.8 | 0.0 |
| **FUS-LCD**<br>(functional) | 19.7 | 18.0 | 59.1 | 0.0 |
| **Aβ42**<br>(pathogenic) | 14.3 | 4.8 | 7.2 | 35.7 |
| **Tau PHFs**<br>(pathogenic) | 12.3 | 8.2 | 15.1 | 24.4 |

## Notes

### Competing Interest Statement

The authors have declared no competing interest.

